# Parkinson’s disease-linked Parkin mutations impair glutamatergic synaptic transmission and plasticity

**DOI:** 10.1101/373597

**Authors:** Mei Zhu, Giuseppe P. Cortese, Clarissa L. Waites

**Affiliations:** Department of Pathology and Cell Biology, Columbia University Medical Center, New York, NY10032; Department of Psychiatry, Columbia University Medical Center; Department of Neuroscience, Columbia University

**Keywords:** Parkin, AMPAR, NMDAR, GluN1, Homer1, Parkinson’s disease, synapse

## Abstract

Parkinson’s disease (PD)-associated E3 ubiquitin ligase Parkin is enriched at glutamatergic synapses, where it ubiquitinates multiple substrates, suggesting that its mutation/loss-of-function could contribute to the etiology of PD by disrupting excitatory neurotransmission. Here, we evaluate the impact of four common PD-associated Parkin point mutations (T240M, R275W, R334C, G430D) on glutamatergic synaptic function in hippocampal neurons. We find that expression of these point mutants in Parkin-deficient and -null backgrounds alters NMDA and AMPA receptor-mediated currents and cell-surface levels, and prevents the induction of long-term depression. Mechanistically, we demonstrate that Parkin regulates NMDA receptor trafficking through its ubiquitination of GluN1, and that all four mutants are impaired in this ubiquitinating activity. Furthermore, Parkin regulates synaptic AMPA receptor trafficking via its binding and retention of the postsynaptic scaffold Homer1, and all mutants are similarly impaired in this capacity. Our findings demonstrate that pathogenic Parkin mutations disrupt glutamatergic synaptic transmission and plasticity by impeding NMDA and AMPA receptor trafficking, and through these effects likely contribute to the pathophysiology of PD in *PARK2* patients.

## Background

Mutations in the *PARK2* gene are the most common cause of Autosomal Recessive Juvenile Parkinsonism and a major contributor to familial and sporadic early-onset Parkinson’s disease (PD) (1-4). *PARK2* encodes Parkin, a RING-between-RING domain E3 ubiquitin ligase that catalyzes the covalent attachment of ubiquitin to specific substrates and regulates vital cellular processes including mitochondrial quality control and apoptosis (5-7). Although it remains unclear how Parkin loss-of-function precipitates the death of midbrain dopaminergic neurons to cause PD, its ubiquitination of mitochondrial proteins downstream of the kinase PINK1 has been shown to mediate mitophagy, a selective form of autophagy (8-11). The buildup of damaged and dysfunctional mitochondria promotes oxidative stress, to which dopaminergic neurons are particularly vulnerable (12, 13), potentially explaining one mechanism through which *PARK2* mutations induce dopaminergic cell loss and the motor symptoms of PD.

However, Parkin is highly expressed throughout the brain and known to regulate other aspects of neuronal function, including glutamatergic neurotransmission (14). Indeed, defects in glutamatergic transmission and plasticity are reported at hippocampal and corticostriatal synapses deficient in Parkin (15-20). Parkin’s mechanisms of action at excitatory synapses remain poorly understood, although its ubiquitinating activity has been found to regulate the stability and function of multiple synaptic substrates, including the presynaptic vesicle-associated protein synaptotagmins XI and IV, the postsynaptic scaffold PICK1, and the kainate receptor subunit GluK2 (21-26). Furthermore, our recent work demonstrates that Parkin also has a structural role at the synapse, linking postsynaptic endocytic zones required for AMPA-type glutamate receptor (AMPAR) capture and internalization to the postsynaptic density through a direct interaction with the scaffold protein Homer1 (19). In Parkin deficient neurons, both the levels of postsynaptic Homer1 and the density of endocytic zones are significantly reduced, leading to impaired AMPAR retention at synapses and ultimately to decreased AMPAR-mediated currents (19).

Loss of these enzymatic and structural roles of Parkin at glutamatergic synapses likely contributes to the symptoms and progression of PD in patients with *PARK2* mutations. Consistent with this concept, PD is recognized as a multi-system disorder with both motor and non-motor symptoms, including resting tremor, muscle rigidity, disordered sleep, sensory dysfunction, depression, and cognitive impairment (27, 28). Although some of the >200 pathogenic mutations, deletions, and exonic rearrangements identified in *PARK2* have been shown to disrupt Parkin’s E3 ligase activity (29-32), their effects at glutamatergic synapses, comprising the vast majority of synapses in the brain, are almost completely unexplored.

Here, we evaluate the effects of four PD-associated Parkin point mutations (T240M, R275W, R334C, G430D) on neurotransmission and plasticity in hippocampal neurons, which are both rich sources of glutamatergic synapses as well as critical substrates for learning and memory. We find that all four mutants alter NMDA-and AMPA-type glutamate receptor trafficking and signaling. Mechanistically, we identify NMDA receptor (NMDAR) subunit GluN1 as a novel Parkin substrate, and find that the mutants are defective in GluN1 ubiquitination, leading to decreased cell-surface NMDAR levels. Furthermore, the mutants exhibit reduced binding and synaptic retention of Homer1, leading to decreased cell-surface AMPAR levels and impaired AMPAR internalization during the induction of synaptic depression. Taken together, these data demonstrate that pathogenic Parkin mutations impair glutamatergic neurotransmission and plasticity, likely contributing to the etiology of *PARK2*-associated PD.

## Results

### Pathogenic Parkin mutations alter AMPAR and NMDAR cell-surface levels and currents

More than 200 pathogenic mutations have been identified in *PARK2*, comprising point mutations, deletions, and exonic rearrangements that are associated with early-onset PD and autosomal recessive juvenile PD (5, 35, 36). However, few of these have been evaluated for their impact on excitatory synapse function. We examined four of the most prevalent Parkin point mutations, collectively identified in ~100 families worldwide (Parkinson Disease Mutation Database; http://www.molgen.vib-ua.be/PDMutDB) and located across three domains of Parkin (T240M and R275W in the RING1 domain; R334C in the IBR domain; G430D in the RING2 domain)(3, 37-39)(Additional file 1: Figure S1a). We first assessed the expression and subcellular localization of these mutants in cultured hippocampal neurons from either the Parkin knockdown background (*i.e.* lentivirally transduced on 2 days *in vitro* (DIV) with an shRNA against rat Parkin (shParkin), previously shown to reduce Parkin levels by ~50% (19)), or Parkin knockout background (*i.e.* cultured from KO rats; SAGE/Horizon Discovery). GFP-tagged T240M, R275W, R334C, and G430D human Parkin constructs exhibited similar subcellular localizations to GFP-wild-type (WT) Parkin, and three of these mutants were expressed at similar levels (Fig. 1; Additional file 1: Figure S1b-d). G430D expression was somewhat higher in the knockdown background (Fig. S1b, d), perhaps reflecting a deficiency in Parkin auto-ubiquitination and degradation for this mutant.

**Figure 1:**
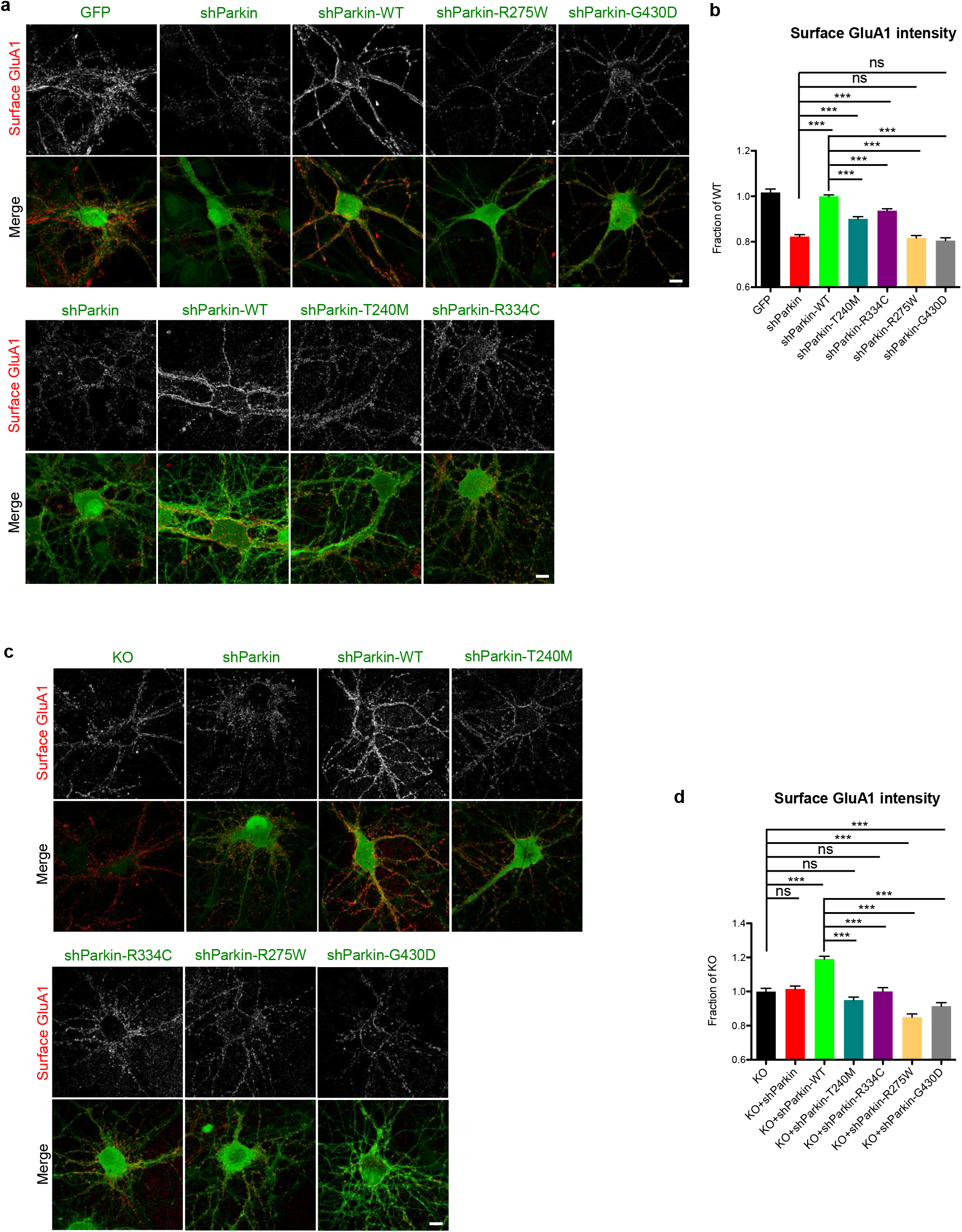
Parkin mutation/loss-of-function decreases cell-surface GluA1 levels. (a) Representative images of surface GluA1 staining (red) in 14-16 DIV hippocampal neurons expressing GFP, shParkin, shParkin-WT, shParkin-T240M, shParkin-R275W, shParkin-R334C or shParkin-G430D constructs. Scale bar, 10 μm. (b) Quantification of cell-surface GluA1 intensity expressed as a fraction of shParkin-WT (n ≥70 fields of view per condition with >100 GluA1 puncta per field, results confirmed in 4 independent experiments. \*\*\**P*<0.001, one-way ANOVA, error bars represent SEM). (c) Representative images of surface GluA1 staining (red) in 14-16 DIV Parkin KO hippocampal neurons expressing shParkin, shParkin-WT, shParkin-T240M, shParkin-R275W, shParkin-R334C or shParkin-G430D constructs and non-transduced KO control. Scale bar, 10 μm. (d) Quantification of cell-surface GluA1 intensity expressed as a fraction of Parkin KO (n ≥50 fields of view per condition with >100 GluA1 puncta per field, results confirmed in 2 independent experiments. \*\*\**P*<0.001, one-way ANOVA, error bars represent SEM).

We previously reported that Parkin knockdown/knockout decreased cell-surface AMPA receptor (AMPAR) levels, and that co-expression of WT-Parkin (shParkin-WT) rescued this phenotype to the level of GFP-expressing control neurons (19). We therefore assessed whether expression of each of the four Parkin mutants could similarly rescue cell-surface AMPAR levels. As in our previous study, we observed that shParkin decreased GluA1 levels by nearly 20% compared to GFP or shParkin-WT expression (Fig. 1a, b). None of the four mutants could match WT Parkin restoration of GluA1 levels, although both T240M and R334C were able to partially rescue this phenotype (Fig. 1a, b). To rule out the ability of the mutants to act as dominant-negatives on endogenous Parkin, we also examined their effects in the KO background. Here, the mutants were similarly unable to increase surface GluA1 levels above those observed in the KO condition, while WT Parkin expression increased GluA1 levels by 20% (Fig. 1c, d), equivalent to its level of rescue in knockdown neurons (Fig. 1a, b). Moreover, expression of shParkin in the KO neurons did not further reduce or alter GluA1 levels (Fig. 1c, d), indicating its lack of off-target effects.

Given Parkin’s reported ability to regulate other glutamate receptor types (*e.g*. kainate) via ubiquitination (26), we also examined cell-surface NMDA receptor levels in these neurons by immunostaining with a human-derived antibody against the GluN1 subunit (40). Intriguingly, we observed that shParkin similarly reduced cell-surface GluN1 levels (by ~15%) compared to GFP or shParkin-WT expression (Fig. 2a, b). Again, three of the mutants (R334C, R275W, G430D) were unable to rescue this phenotype, while T240M rescued to the level of WT Parkin (Fig. 2a, b). These experiments were also performed in Parkin KO neurons, where it was confirmed that none of the mutants, including T240M, was able to increase surface GluN1 levels above those observed in the KO or KO + shParkin conditions (Fig. 2c, d). In contrast, WT Parkin expression increased GluN1 levels by ~20%, similar to its level of rescue in shParkin-expressing neurons (Fig. 2c, d). Together, these findings indicate that none of the four Parkin mutants are able to support normal cell-surface AMPAR and NMDAR levels.

**Figure 2:**
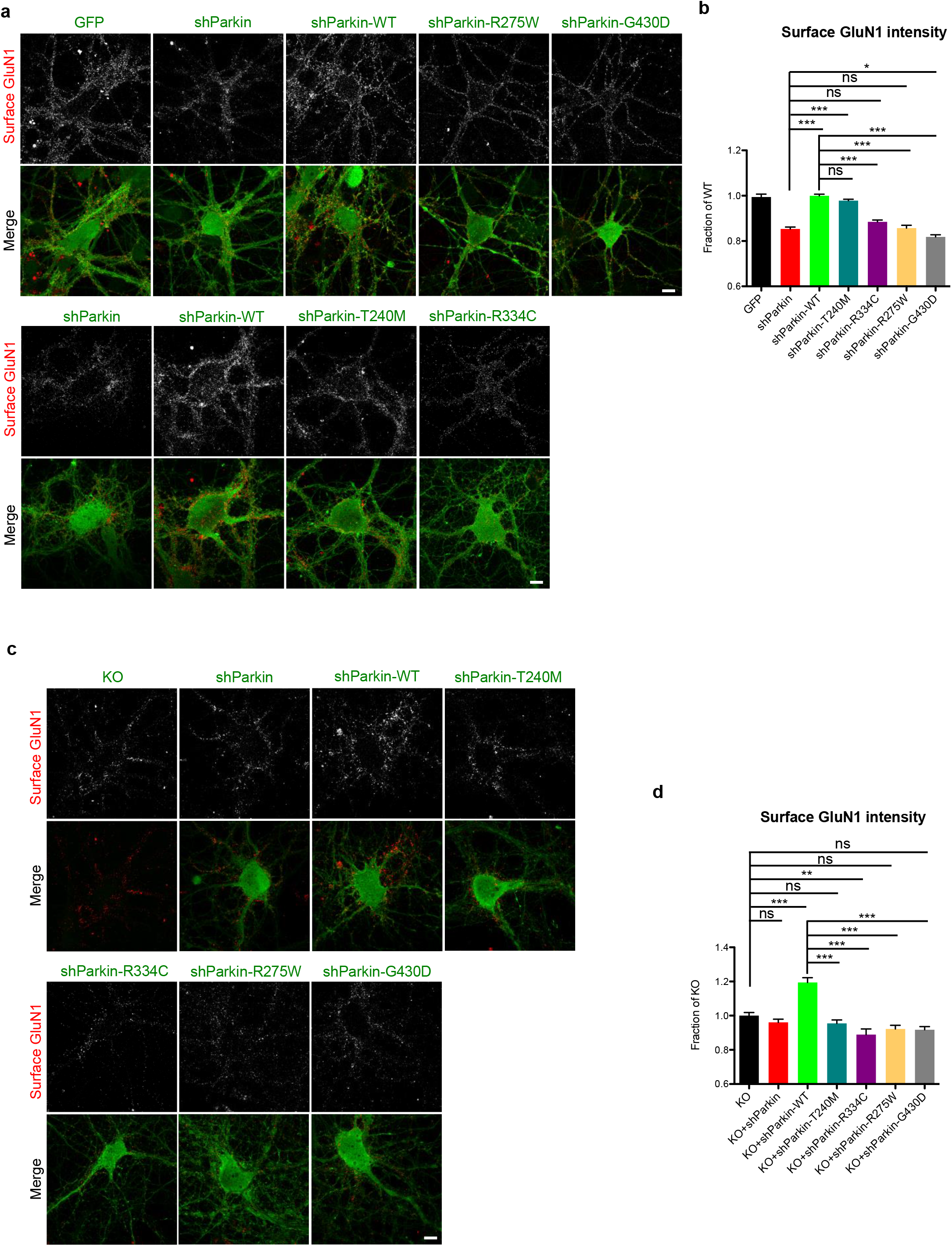
Parkin mutation/loss-of-function decreases cell-surface GluN1 levels. (a) Representative images of surface GluN1 staining (red) in 14-16 DIV hippocampal neurons expressing GFP, shParkin, shParkin-WT, shParkin-T240M, shParkin-R275W, shParkin-R334C or shParkin-G430D constructs. Scale bar, 10 μm. (b) Quantification of cell-surface GluN1 intensity expressed as a fraction of shParkin-WT (n ≥50 fields of view per condition with >100 GluN1 puncta per field, results confirmed in 4 independent experiments. \*\*\**P*<0.001, one-way ANOVA, error bars represent SEM). Scale bar, 10 μm. (c) Representative images of surface GluN1 staining (red) in 14-16 DIV Parkin KO hippocampal neurons expressing shParkin, shParkin-WT, shParkin-T240M, shParkin-R275W, shParkin-R334C or shParkin-G430D constructs, and non-transduced KO control. Scale bar, 10 μm. (d) Quantification of cell-surface GluN1 intensity expressed as a fraction of Parkin KO (n ≥50 fields of view per condition with >100 GluA1 puncta per field, results confirmed in 2 independent experiments. \*\*\**P*<0.001, one-way ANOVA, error bars represent SEM).

To determine whether the Parkin mutants compromise excitatory neurotransmission, we examined their effects on AMPA and NMDA receptor spontaneous excitatory postsynaptic currents (EPSCs) in both Parkin knockdown and KO neurons. We compared the degree of rescue from expression of Parkin mutants to that of WT Parkin, which significantly increased AMPAR miniature EPSC (mEPSC) amplitudes in the knockdown and KO backgrounds (Fig. 3a, b, d, e), consistent with our previous findings (19) and results with cell-surface GluA1 immunostaining (Fig. 1). WT Parkin expression also increased mEPSC frequency in the KO but not knockdown background (Fig. 3a, c, d, f), as expected given our previous results showing that AMPAR mEPSC frequency was decreased in Parkin KO but not knockdown neurons (19). In contrast, none of the Parkin mutants was able to increase mEPSC amplitude above knockdown or KO levels, and none rescued mEPSC frequency in KO neurons, although T240M and R275W did have small but significant effects on frequency in the knockdown background (Fig. 3a-f).

**Figure 3:**
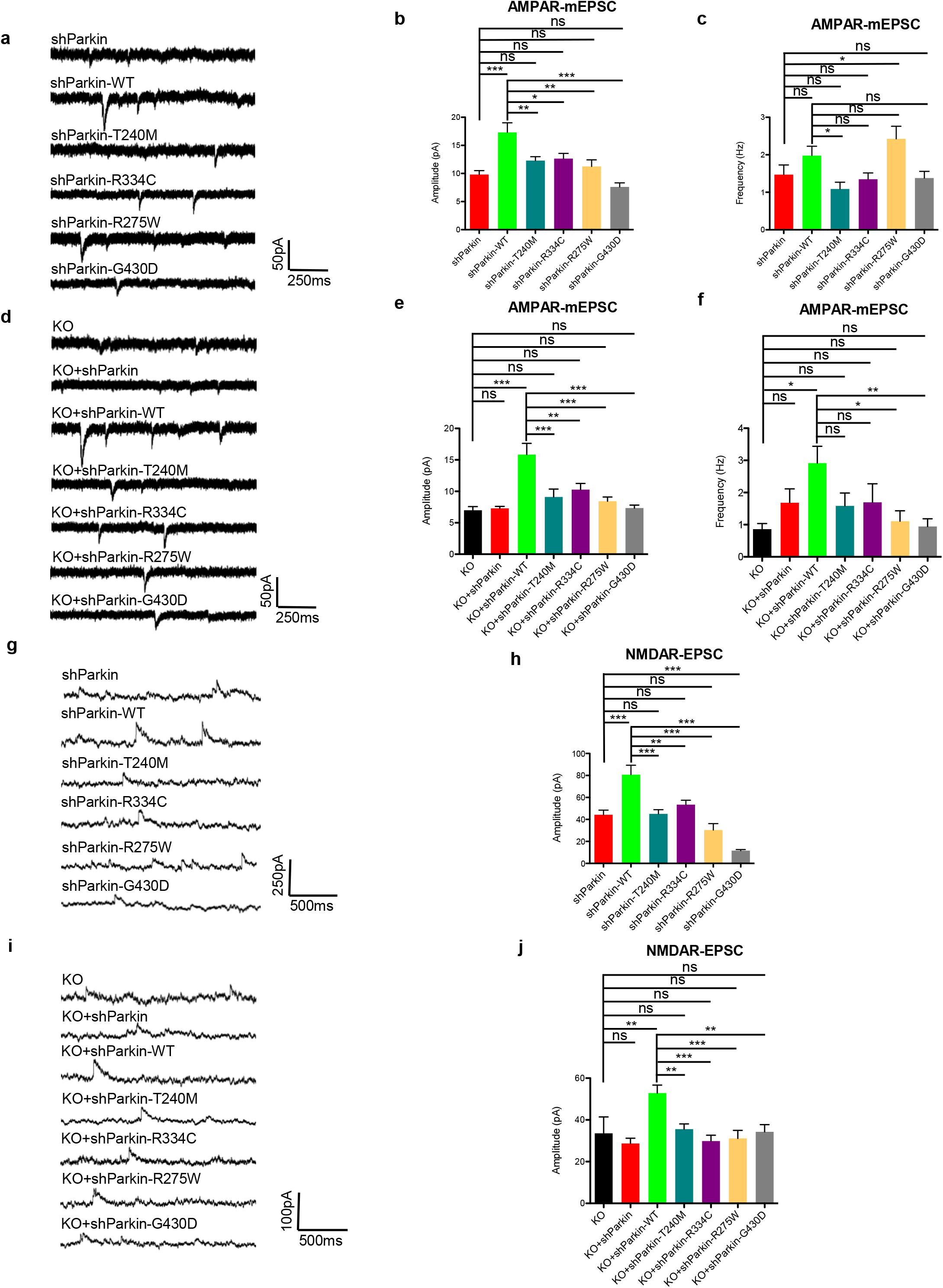
Parkin mutation/loss-of-function decreases AMPA and NMDA receptor-mediated currents. (a) Representative traces of spontaneous AMPAR mEPSCs from 14-16 DIV hippocampal neurons transduced with shParkin +/- WT Parkin or T240M/R275W/R334C/G430D mutants. (b, c) Quantification of mEPSC amplitude (b) and frequency (c) for the six conditions (n=14 for shParkin, shParkin-WT and shParkin-T240M, 10 for shParkin-R334C, 8 for shParkin-R275W and shParkin-G430D; \**P*<0.05, \*\**P*<0.005, \*\*\**P*<0.001, one-way ANOVA, error bars represent SEM). (d) Representative traces of spontaneous AMPAR mEPSCs from 14-16 DIV Parkin KO hippocampal neurons transduced with shParkin +/- WT Parkin or T240M/R275W/R334C/G430D mutants and non-transduced KO control. (e, f) Quantification of mEPSC amplitude (e) and frequency (f) for the seven conditions (n=12 for shParkin-WT and shParkin-T240M, 11 for shParkin-R275W and shParkin-G430D, 10 for shParkin-R334C, 8 for shParkin and non-transduced KO control; \**P*<0.05, \*\**P*<0.005, \*\*\**P*<0.001, one-way ANOVA, error bars represent SEM). (g) Representative traces of NMDAR currents (NMDAR-EPSC) from 14-16 DIV hippocampal neurons transduced with shParkin +/- WT Parkin or T240M/R275W/R334C/G430D mutants. (h) Quantification of NMDAR-EPSC amplitude for the six conditions (n=12 for shParkin-T240M and shParkin-R334C, 8 for shParkin, shParkin-WT, shParkin-R275W and shParkin-G430D; \*\**P*<0.005, \*\*\**P*<0.001, one-way ANOVA, error bars represent SEM). (i) Representative traces of NMDAR currents (NMDA-EPSC) from 14-16 DIV Parkin KO hippocampal neurons transduced with shParkin +/- WT Parkin or T240M/R275W/R334C/G430D mutants and non-transduced KO control. (j) Quantification of NMDAR-EPSC amplitude for the seven conditions (n=12 for shParkin-WT and shParkin-T240M, 11 for shParkin-G430D, 10 for shParkin-R334C and shParkin-R275W, 8 for shParkin and non-transduced KO control; \*\**P*<0.005, \*\*\**P*<0.001, one-way ANOVA, error bars represent SEM).

We also examined spontaneous EPSC amplitudes for NMDARs (NMDAR-EPSCs) in both Parkin knockdown and KO neurons. Here, we observed that WT Parkin expression significantly increased NMDAR-EPSCs in both of these backgrounds (Fig. 3g-j), indicative of the reduction in synaptic NMDAR levels in the absence of Parkin. Again, the four Parkin mutants were unable to recapitulate WT Parkin function and did not increase NMDAR-EPSC amplitudes above knockdown/KO levels (Fig. 3g-j). To further confirm this loss of NMDARs in the Parkin deficient background, we measured whole cell NMDAR currents in response to local application of NMDA for neurons expressing soluble GFP, human Parkin (Hu-Parkin), shParkin, or shParkin-WT (Additional file 2: Fig. S2a), as well as the levels of NMDAR subunits (GluN1, GluN2A, and GluN2B) by cell-surface biotinylation (Additional file 2: Fig. S2b and S2c). We observed that shParkin significantly reduced both NMDAR currents and surface GluN subunit biotinylation (Additional file 2: Fig. S2), confirming by several methods that Parkin regulates cell-surface NMDAR levels.

### Parkin’s E3 ligase activity is required for maintaining surface NMDAR but not AMPAR levels

We next investigated the mechanisms through which these pathogenic mutations disrupt cell-surface AMPAR and NMDAR expression. Each mutant is reported or predicted to impair Parkin’s E3 ligase activity (29, 30, 32), suggesting that the loss of Parkin-mediated ubiquitination leads to a reduction of cell-surface glutamate receptor levels. To test this, we took advantage of two mutants engineered to augment or inhibit Parkin-mediated ubiquitination. In particular, mutation of tryptophan 403 to alanine (W403A) was shown to increase Parkin’s E3 ligase activity, and mutation of cysteine 431 to serine (C431S) to inhibit this activity (29)(Additional file 1: Fig S1a). As in previous experiments, we expressed WT, W403A, or C431S Parkin together with shParkin in hippocampal neurons, and after verifying their expression (Additional file 1: Fig. S1e), immunostained with GluN1 or GluA1 antibodies. Unexpectedly, we found that both W403A and C431S mutants efficiently restored surface GluA1 to WT Parkin levels (Fig. 4a, c), while only W403A rescued surface GluN1 levels (Fig. 4b, d). These findings demonstrate that Parkin’s E3 ligase activity is essential for maintaining cell-surface NMDAR but not AMPAR levels, and indicate that Parkin regulates each receptor type through a distinct mechanism. Further supporting this concept, we found that overexpression of the postsynaptic scaffold Homer1, previously shown to rescue surface GluA1 levels in Parkin deficient neurons, did not rescue surface GluN1 levels (Additional file 3: Fig. S3).

**Figure 4:**
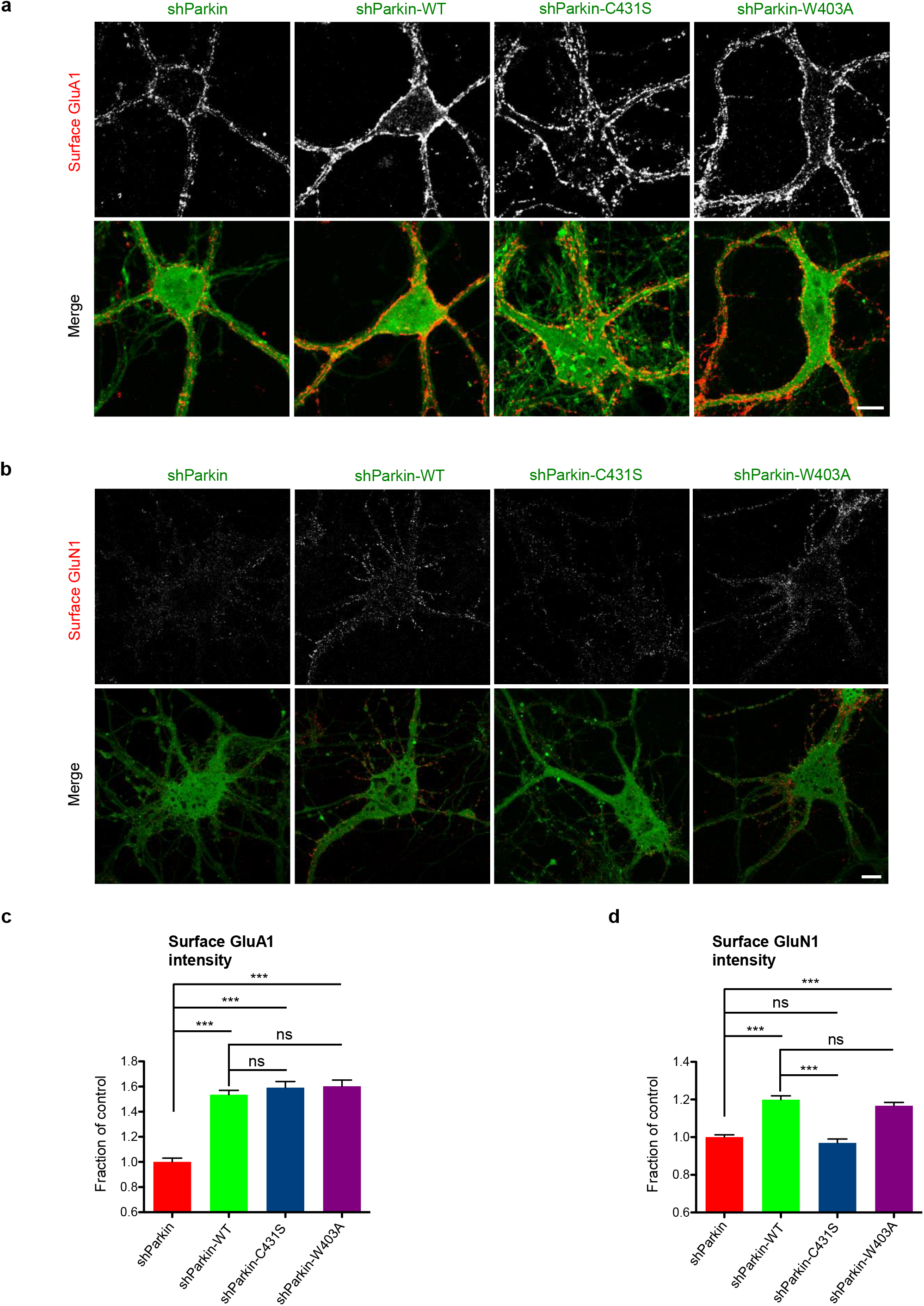
Parkin E3 ligase activity is required to maintain cell-surface NMDARs but not AMPARs. (a) Representative images of surface GluA1 staining (red) in 14-16 DIV hippocampal neurons expressing shParkin +/- WT, C431S, or W403A Parkin constructs. Scale bar, 10 μm. (b) Same condition as (a), but for surface GluN1 staining (red). Scale bar, 10 μm. (c) Quantification of cell-surface GluA1 intensity expressed as a fraction of shParkin control (n ≥40 fields of view per condition with >100 GluA1 puncta per field, results confirmed in 3 independent experiments. \*\*\**P*<0.001, one-way ANOVA, error bars represent SEM). (d) Quantification of cell-surface GluN1 intensity expressed as a fraction of shParkin control (n≥40 fields of view per condition with >100 GluN1 puncta per field, results confirmed in 3 independent experiments. \*\*\**P*<0.001, one-way ANOVA, error bars represent SEM).

### Parkin mutants are deficient in ubiquitination of NMDAR subunit GluN1

To determine whether Parkin regulation of surface NMDAR levels is mediated through direct ubiquitination of NMDAR subunits, we performed ubiquitination assays in HEK293T cells co-transfected with GFP-tagged NMDAR or AMPAR subunits (GluN1, GluN2B, GluA1, GluA2) together with HA-ubiquitin, Myc vector control or Myc-Parkin. Here, we found that Parkin co-expression significantly increased ubiquitination of GFP-GluN1 (Fig. 5a, b), and that a similar degree of GluN1 ubiquitination was observed following its immunoprecipitation under denaturing conditions (1% SDS) to eliminate other potential binding partners (Additional file 4: Fig. S4a, b). In contrast, the other glutamate receptor subunits had ubiquitination levels equivalent to that of GFP control (Fig. 5a, b). Follow-up co-immunoprecipitation assays revealed that GluN1 was also the only subunit to co-precipitate with Myc-Parkin (Fig. 5c). These data demonstrate that Parkin specifically interacts with and ubiquitinates the GluN1 subunit of NMDARs.

**Figure 5:**
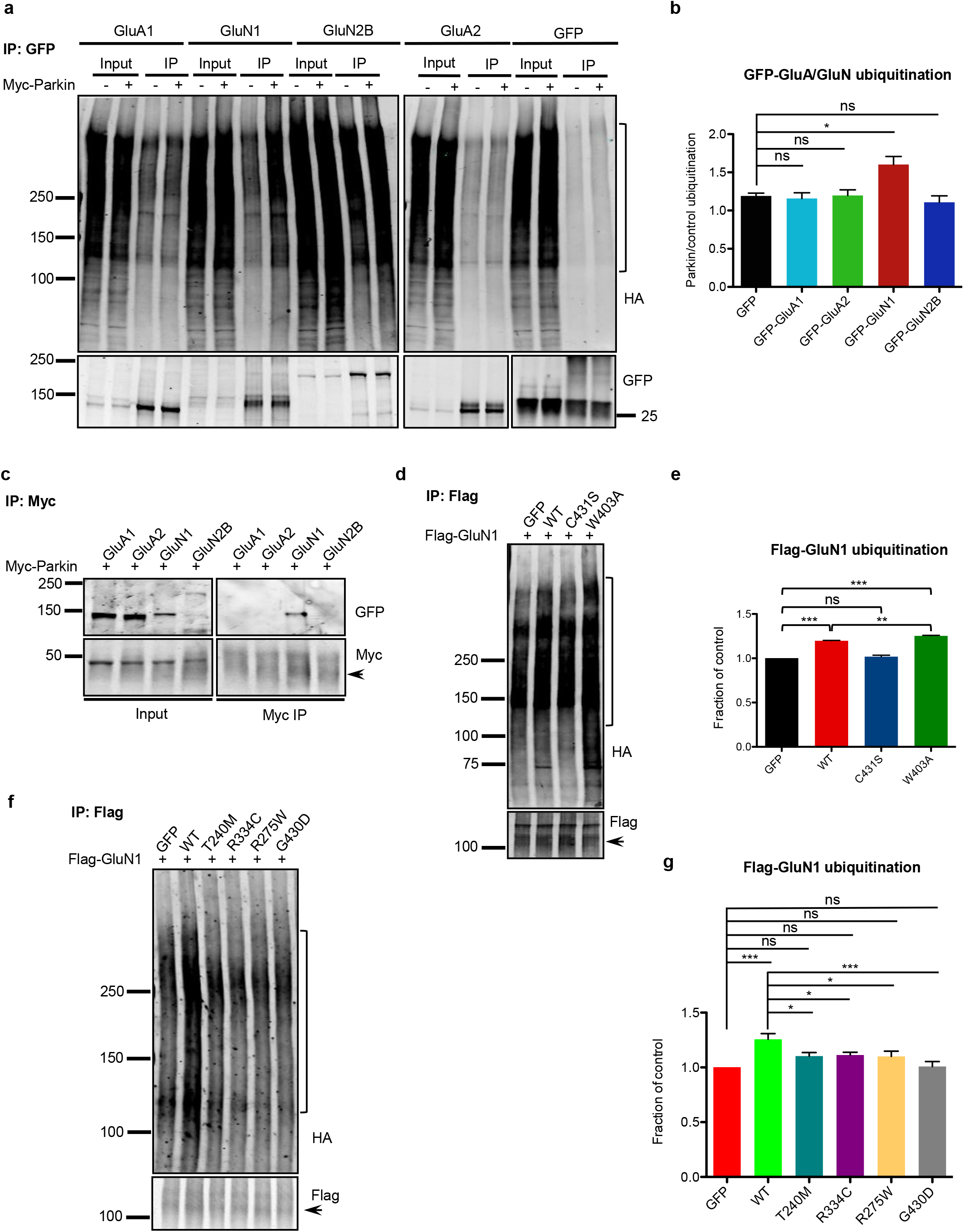
Parkin-mediated GluN1 ubiquitination is impaired in pathogenic mutants. (a) Representative immunoblots for GFP immunoprecipitation (IP) from HEK293T cell lysates expressing Myc/Myc-Parkin, GFP-GluA1/-GluA2/-GluN1/-GluN2B, and HA-ubiquitin, probed for HA and GFP. Ubiquitin immunoreactivity used for quantification is marked on HA blots. (b) Quantification of GFP-GluA or GluN ubiquitination, expressed as the ratio of marked HA blot intensity (a) with Myc-Parkin (+) to Myc control (-), then normalized to immunoprecipitated GFP, GFP-GluA or GluN (n=3 experiments, \**P*<0.05; one-way ANOVA, error bars represent SEM). (c) Representative Myc and GFP immunoblots for Myc IP from HEK293T cell lysates expressing Myc-Parkin and GFP-GluA1/-GluA2/-GluN1/-GluN2B. Arrowhead indicates immunoprecipitated Myc-Parkin (just below IgG band). (d) Representative HA and Flag immunoblots for Flag IP from HEK293T cell lysates expressing Flag-GluN1, GFP control/GFP-Parkin WT/C431S/W403A, and HA-ubiquitin. Arrowhead indicates immunoprecipitated Flag-GluN1. Ubiquitin immunoreactivity used for quantification is marked on HA blots. (e) Quantification of Flag-GluN1 ubiquitination by measurement of marked HA blot intensity (d), normalized to immunoprecipitated Flag-GluN1 and reported as a fraction of GFP control (n=3 experiments, \*\**P*<0.01, \*\*\**P*<0.001, one-way ANOVA, error bars represent SEM). (f) Representative HA and Flag immunoblots for Flag IP from HEK293T cell lysates expressing Flag-GluN1, GFP control/GFP-Parkin WT/T240M/R275W/R334C/G430D constructs, and HA-ubiquitin. Arrowhead indicates immunoprecipitated Flag-GluN1. Ubiquitin immunoreactivity used for quantification is marked on HA blots. (g) Quantification of Flag-GluN1 ubiquitination by measurement of marked HA intensity (f), normalized to immunoprecipitated Flag-GluN1 and reported as a fraction of GFP condition (n=3 experiments, \**P*<0.05; \*\**P*<0.01, \*\*\**P*<0.001, one-way ANOVA, error bars represent SEM).

To evaluate whether the GFP-tagged Parkin mutants were impaired in their ability to ubiquitinate GluN1, we performed ubiquitination assays using Flag-GluN1. We first tested the sensitivity of the assay by examining GluN1 ubiquitination by the W403A (active) and C431S (inactive) Parkin mutants. Here, we found that W403A increased GluN1 ubiquitination compared to WT Parkin, while ubiquitination by C431S was indistinguishable from soluble GFP control (Fig. 5d, e). We next examined Flag-GluN1 ubiquitination in the presence of T240M, R275W, R334C, and G430D, and found the levels of ubiquitin immunoreactivity to be similar to those seen in the control condition (Fig. 5f, g), confirming that these Parkin mutants are deficient in their ubiquitination of GluN1.

### Parkin regulates NMDAR internalization and recycling

Ubiquitination is known to mediate protein degradation, and Parkin-mediated ubiquitination regulates the stability and degradation of multiple synaptic substrates (21-24, 26). However, since cell-surface GluN1 levels were decreased rather than increased following Parkin loss-of-function, we hypothesized that Parkin-mediated ubiquitination does not regulate NMDAR degradation. This hypothesis was confirmed by cycloheximide-chase experiments to compare the degradation of NMDAR subunits (GluN1, GluN2A, GluN2B) in control neurons with those expressing shParkin +/- WT Parkin (Additional file 4: Fig. S4c, d). Since ubiquitination also regulates glutamate receptor trafficking (41-43), we utilized antibody feeding assays to monitor the internalization and recycling of GluN1 in neurons expressing soluble GFP or shParkin +/- WT-Parkin (see Methods for details). Here, we found that both the internalization of GluN1 and its recycling back to the cell-surface were significantly impaired in Parkin knockdown neurons, and that these defects were fully rescued by WT Parkin (Fig. 6a-d). Altogether, these findings indicate that Parkin-mediated ubiquitination regulates both the endocytosis and exocytosis of NMDARs at the cell-surface, leading to an overall reduction of surface NMDARs in Parkin deficient neurons.

**Figure 6:**
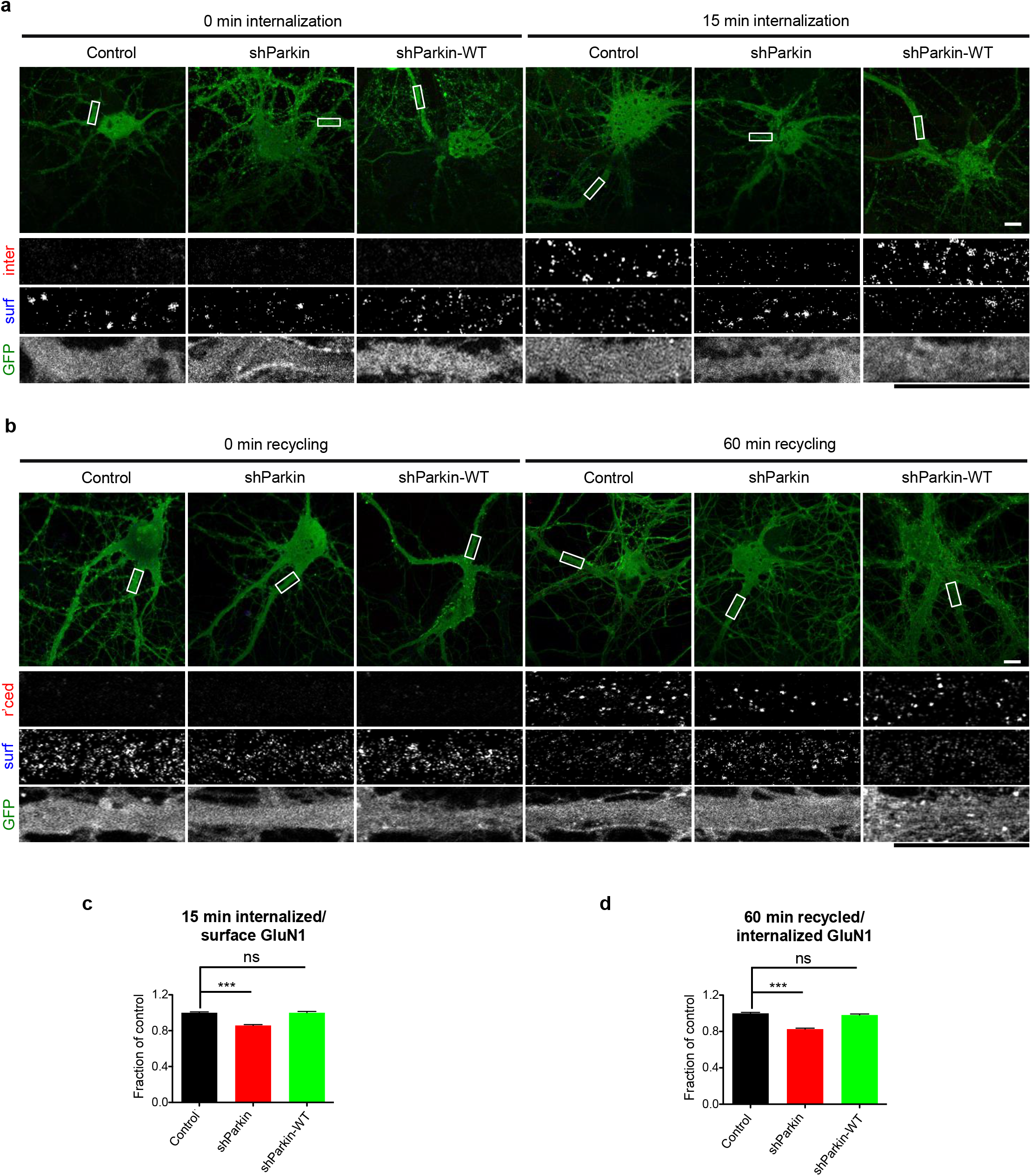
Parkin deficiency impairs GluN1 internalization and recycling. (a) Representative images of 14-16 DIV hippocampal neurons expressing soluble GFP control/shParkin/shParkin-WT constructs, immunostained for cell surface (surf) and internalized (inter) GluN1 after 0 and 15min internalization at 37°C. Scale bars, 10 μm. (b) Representative images of 14-16 DIV hippocampal neurons expressing the same constructs as in (a), immunostained for cell surface (surf) and recycled (rec’d) GluN1 after 30 min internalization followed by 0 and 60 min recycling at 37°C. Scale bars, 10 μm. (c) Quantification of GluN1 internalization at 15 min, expressed as the ratio of internalized to cell surface GluN1 and normalized to GFP control condition (n ≥50 fields of view per condition with >50 GluN1 puncta per field, results confirmed in 3 independent experiments; \*\*\**P*<0.001, unpaired *t* test, error bars represent SEM). (d) Quantification of GluN1 recycling at 60 min, expressed as the ratio of recycled to internalized GluN1 and normalized to GFP control condition (n ≥55 fields of view per condition with >50 GluN1 puncta per field, results confirmed in 3 independent experiments; \*\*\**P*<0.001, unpaired *t* test, error bars represent SEM).

### Parkin mutants are impaired in their binding and retention of postsynaptic Homer1

Although defective GluN1 ubiquitination provides a mechanistic explanation for how Parkin mutants decrease cell-surface NMDAR levels, it does not explain how these mutants decrease cell-surface AMPAR levels, a phenotype that does not appear to depend on Parkin’s ubiquitinating activity (see Fig. 4). We previously showed that direct binding of Parkin to Homer1 is necessary for maintaining surface AMPAR levels, by tethering endocytic zones (EZs) for AMPAR capture and internalization to the postsynaptic density (19). In Parkin deficient neurons, Homer1 levels are decreased, leading to reduced EZ density and loss of synaptic AMPARs (19). To determine whether the Parkin mutants were impaired in their binding to Homer1, we performed co-immunoprecipitation assays in HEK293T cells co-expressing GFP-tagged Homer1 and either T240M, R275W, R334C, or G430D. Intriguingly, all of the mutants exhibited significantly less co-precipitation with Homer1 than did WT Parkin (Fig. 7a, b), reinforcing the link between Parkin/Homer1 binding and surface GluA1 levels. We also performed quantitative immunofluorescence microscopy to measure synaptic Homer1 levels in knockdown neurons expressing WT Parkin or each mutant. Here, we found that synaptic Homer1 levels were significantly decreased in the presence of all four mutants compared to WT Parkin (Fig. 7c, d), consistent with our immunostaining and electrophysiology data showing decreased cell-surface AMPARs and mEPSC amplitudes (Figs. 1 and 3). Together, these findings provide an explanation for how Parkin mutations disrupt synaptic AMPAR levels and AMPAR-mediated currents.

**Figure 7:**
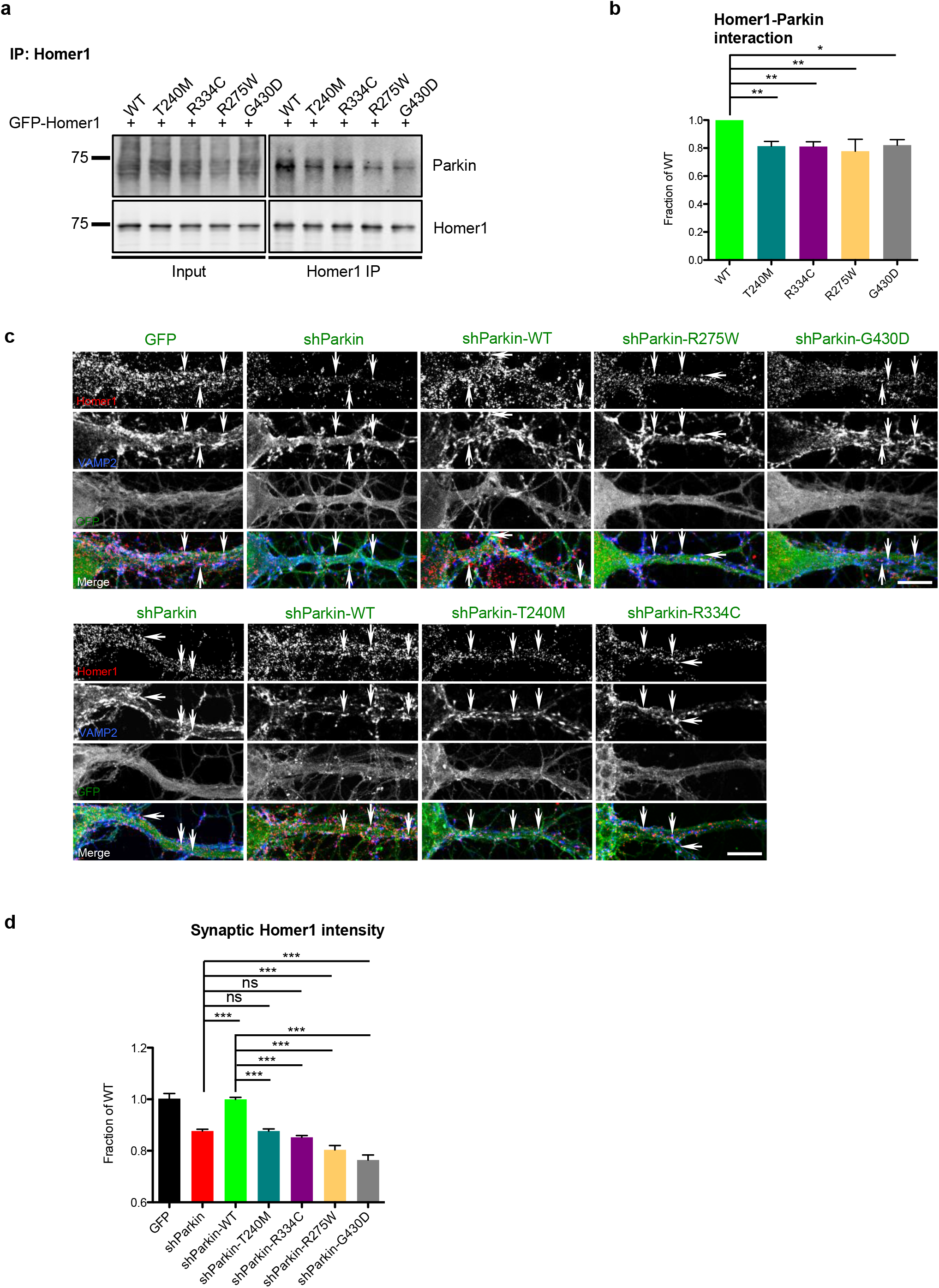
Parkin mutations reduce Homer1 binding and synaptic retention. (a) Representative Parkin and Homer1 immunoblots for Homer1 IP from HEK293T cell lysates expressing GFP-Homer1 and GFP-Parkin WT/T240M/R275W/R334C/G430D. (b) Quantification of Homer1/Parkin interaction by measurement of co-immunoprecipitated GFP-Parkin WT/T240M/R275W/R334C/G430D, normalized to immunoprecipitated GFP-Homer1 and reported as a fraction of GFP-Parkin WT condition (n=3 experiments, one-way ANOVA, \**P*<0.05, \*\**P*<0.01, error bars represent SEM). (c) Representative images of Homer1 (red) and VAMP2 (blue) staining in 14-16 DIV hippocampal neurons expressing GFP, shParkin, or shParkin-WT/-T240M/-R275W/-R334C/-G430D. Arrows indicate representative synaptic Homer1 puncta (based on co-localization with VAMP2) for each condition. Scale bar, 10 μm. (d) Quantification of synaptic Homer1 intensity at VAMP2-immunopositive puncta along dendrites, expressed as a fraction of shParkin-WT control condition. (n≥50 fields of view per condition with >50 co-localized Homer1 puncta per field, results confirmed in 3 independent experiments; \*\*\**P*<0.001, one-way ANOVA, error bars represent SEM).

### Parkin-mediated AMPAR internalization is required for induction of synaptic depression

NMDAR-dependent synaptic plasticity mechanisms, including long-term potentiation (LTP) and long-term depression (LTD), are responsible for dynamic changes in synaptic strength that mediate important high-order brain functions such as learning and memory, pain perception, addictive behaviors, and mood regulation (44-46). Induction of LTP and LTD require rapid, calcium-mediated changes in the number, phosphorylation state, and/or subunit composition of synaptic cell-surface AMPARs (47, 48). Since we identified clear roles for Parkin in AMPAR and NMDAR trafficking, we hypothesized that Parkin mutation or loss-of-function would impair these synaptic plasticity mechanisms. Indeed, a previous study reported that Parkin KO animals exhibit reduced hippocampal LTP (20). To test this possibility, we first examined NMDAR-dependent synaptic potentiation and depression in hippocampal neurons expressing soluble GFP or shParkin +/- WT Parkin, using chemical stimuli to induce LTP or LTD (cLTP or cLTD; see Methods and (49, 50)). Following these treatments, we calculated the ratio of mean surface GluA1 intensity following cLTP or cLTD induction to that in the untreated control condition, with values >1 representing insertion of GluA1-containing AMPARs at the cell surface (*i.e.* successful cLTP), and values <1 representing internalization of AMPARs from the surface (*i.e.* successful cLTD). Surprisingly, we found that the magnitude of cLTP was nearly identical for GFP control and shParkin-expressing neurons, although both the basal and potentiated levels of surface GluA1 were lower in knockdown neurons than in control neurons (Fig. 8a, b). On the other hand, co-expression of WT Parkin led to a slight impairment of cLTP induction, likely due to occlusion from higher initial surface levels of GluA1 (Fig. 8a, b). In contrast, Parkin knockdown completely prevented the induction of cLTD, and this phenotype was fully restored by WT Parkin expression (Fig. 8c, d). These data are consistent with our previous findings that Parkin regulates AMPAR endocytosis but not exocytosis (19).

**Figure 8:**
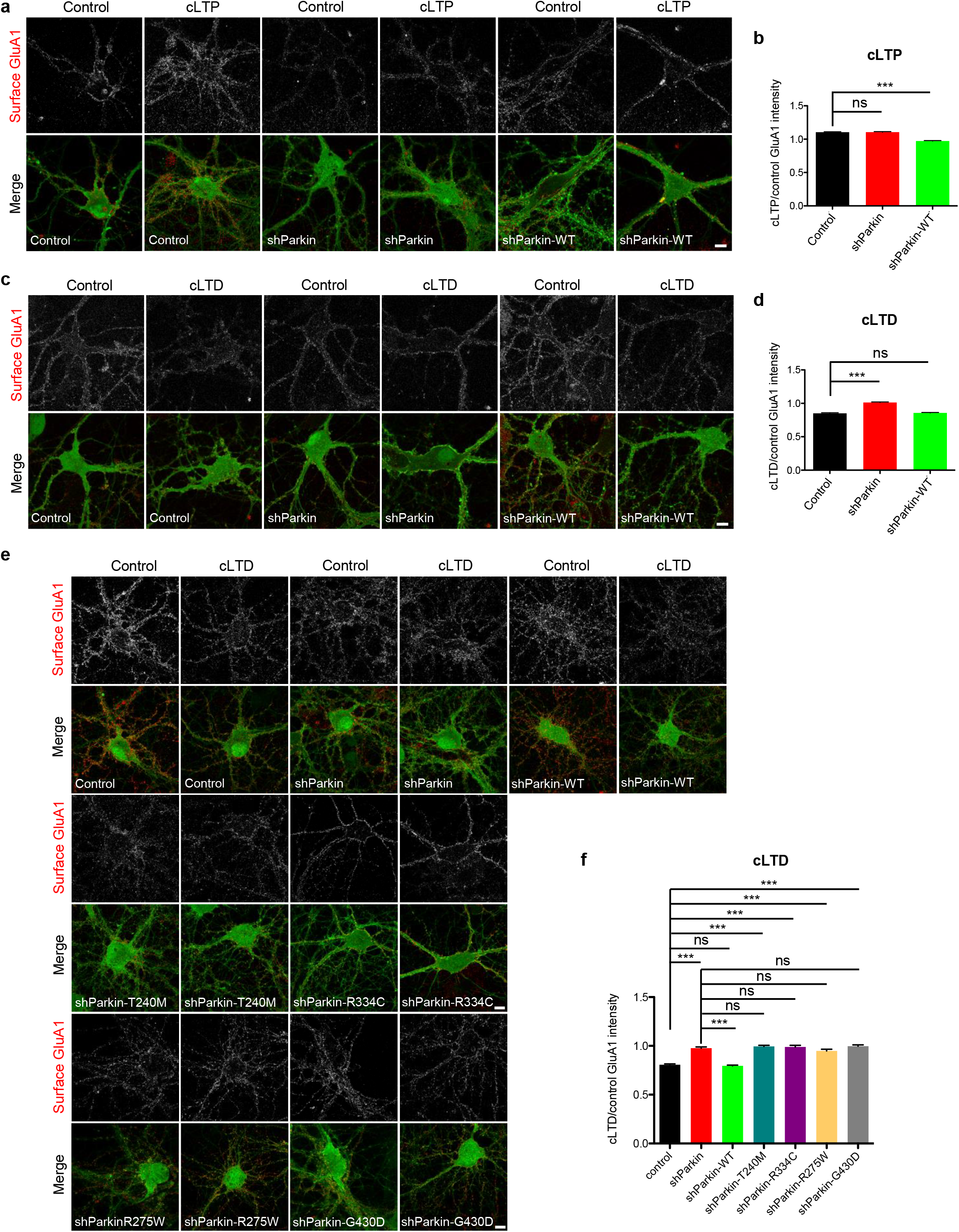
Parkin mutation/knockdown impairs the induction of LTD. (a) Representative images of surface GluA1 staining (red) in 14-16 DIV hippocampal neurons expressing GFP, shParkin or shParkin-WT, under the control condition (no treatment) or after chemical LTP (cLTP) induction. Scale bar, 10 μm. (b) Quantification of the ratio of GluA1 intensity after cLTP induction to the control condition for neurons expressing GFP, shParkin, or shParkin-WT. (c) Same as (a), but for control condition or chemical LTD (cLTD) induction. Scale bar, 10 μm. (d) Quantification of the ratio of GluA1 intensity after cLTD induction to the control condition for neurons expressing GFP, shParkin, or shParkin-WT. (e) Representative images of surface GluA1 staining (red) in 14-16 DIV hippocampal neurons expressing GFP, shParkin or shParkin-WT/-T240M/-R275W/-R334C/-G430D constructs, under the control condition or after cLTD induction. Scale bar, 10 μm. (f) Quantification of the ratio of GluA1 intensity after cLTD induction to the control condition for neurons expressing GFP, shParkin or shParkin-WT/-T240M/-R275W/-R334C/-G430D constructs. For panels (b) and (d), n ≥50 fields of view per condition with >100 GluA1 puncta per field, results confirmed in 3 independent experiments. \*\*\**P*<0.001, unpaired *t* test. For panel (f), n ≥40 fields of view per condition with >100 GluA1 puncta per field, results confirmed in 3 independent experiments. \*\*\**P*<0.001, one-way ANOVA, error bars represent SEM.

We next evaluated whether co-expression of Parkin mutants could rescue this cLTD deficit. Remarkably, we found that none of the four mutants were able to rescue GluA1 internalization following cLTD induction (Fig. 8e, f), consistent with their reduced Homer1 binding and inability to rescue synaptic Homer1 levels compared to WT Parkin (Fig. 7). Supporting the concept that Parkin mutation/loss-of-function prevents LTD induction by disrupting Homer1-linked endocytic zones and thus GluA1 internalization, we found that dephosphorylation of GluA1 at Serine-845, a hallmark of LTD that is dependent upon events upstream of AMPAR internalization (*i.e.* Ca^2+^ influx through NMDARs, phosphatase activation; (49)), was unaffected by Parkin knockdown or mutant co-expression (Additional file 5: Figure S5). These findings indicate that pathogenic Parkin mutations disrupt synaptic depression by impairing AMPAR internalization.

Since hippocampal LTP was reported to be deficient in Parkin KO animals (20), we also examined cLTP induction in hippocampal neurons from the Parkin KO background, in the presence or absence of WT Parkin or the four mutants. We found that the magnitude of cLTP induction as measured by surface GluA1 levels was indistinguishable in KO neurons compared to those expressing WT or mutant Parkin (Fig. 9a, b), again demonstrating that Parkin loss-of-function does not impair LTP. However, induction of cLTD was completely blocked in KO neurons, and this phenotype was rescued only by WT Parkin expression (Fig. 9c, d), indicating that the four mutants cannot support cLTD in either the Parkin deficient or Parkin null backgrounds. Together, these findings indicate that Parkin mediates NMDAR-dependent synaptic depression by facilitating AMPAR internalization, and that pathogenic Parkin mutations prevent this function.

**Figure 9:**
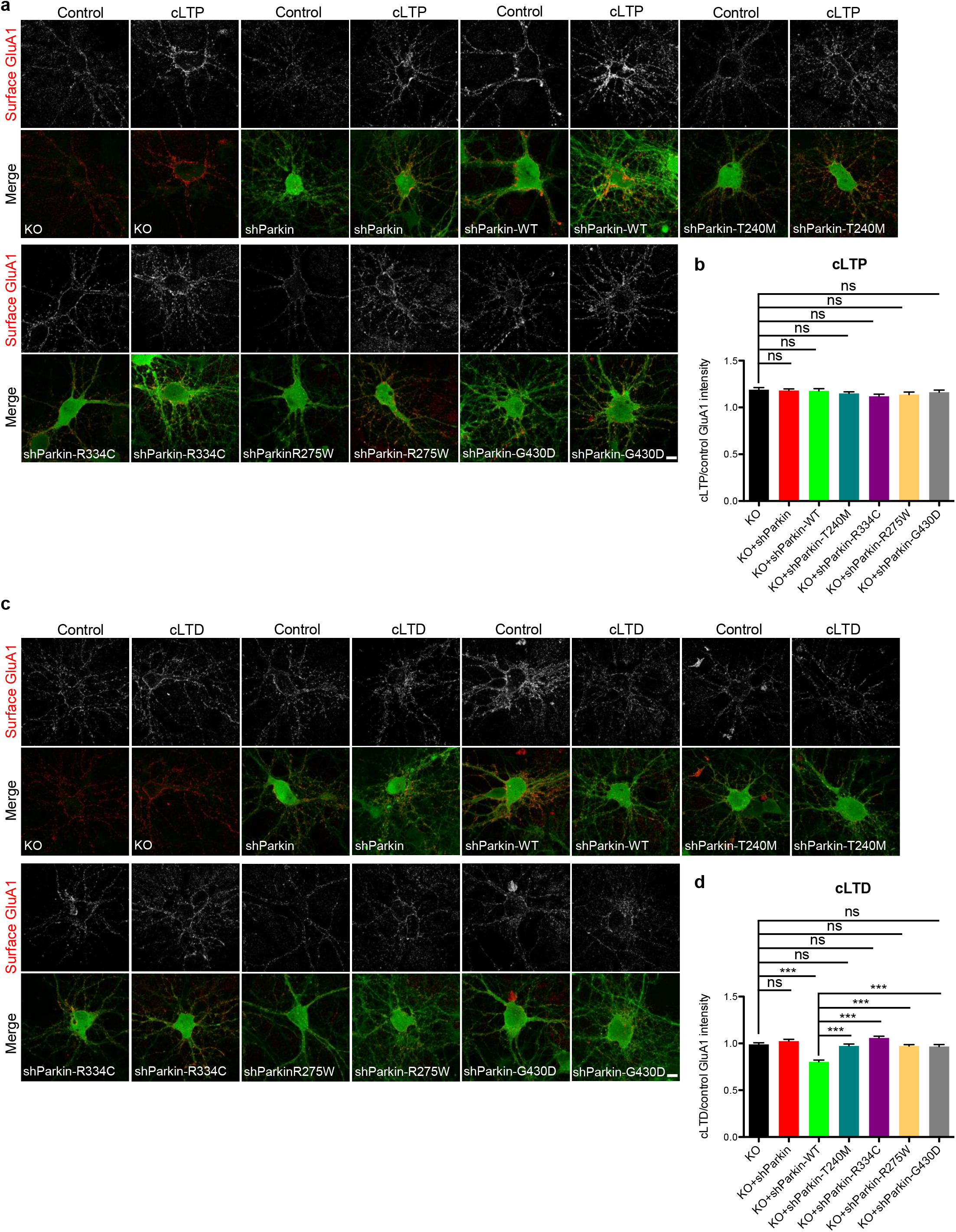
Parkin mutation/knockout impairs the induction of LTD but not LTP. (a) Representative images of surface GluA1 staining (red) in 14-16 DIV Parkin KO hippocampal neurons expressing shParkin or shParkin-WT/-T240M/-R275W/-R334C/-G430D constructs, and non-transduced KO control under the control condition (no treatment) or after chemical LTP (cLTP) induction. Scale bar, 10 μm. (b) Quantification of the ratio of GluA1 intensity after cLTP induction to the control condition for neurons expressing the above Parkin constructs. (c) Same as (a), but for control condition or chemical LTD (cLTD) induction. Scale bar, 10 μm. (d) Quantification of the ratio of GluA1 intensity after cLTD induction to the control condition for neurons expressing above Parkin constructs. For panels (b) and (d), n ≥40 fields of view per condition with >100 GluA1 puncta per field, results confirmed in 2 independent experiments. \*\*\**P*<0.001, one-way ANOVA, error bars represent SEM.

## Discussion

*PARK2* mutations are present in >3% of PD patients, making them more prevalent than mutations in LRRK2 or α-synuclein (38, 39, 51-53). Moreover, PD is a multisystem disorder and Parkin is enriched at postsynaptic densities (PSDs) of glutamatergic synapses, suggesting that *PARK2* mutations could contribute to PD pathophysiology by disrupting excitatory neurotransmission. Here, we demonstrate that expression of four PD-linked Parkin mutations (T240M, R275W, R334C, and G430D) in Parkin-deficient or Parkin null backgrounds, to simulate its heterozygous or homozygous loss-of-function, alters glutamatergic synaptic transmission and plasticity. We provide mechanistic insight into how these Parkin mutations impair NMDAR and AMPAR-mediated signaling, through deficient ubiquitination of GluN1 and deficient binding/retention of postsynaptic Homer1, leading to reduced cell-surface NMDAR and AMPAR levels and impaired AMPAR internalization. Our findings not only demonstrate that Parkin regulates postsynaptic NMDA-and AMPA-type glutamate receptors through distinct mechanisms, but also that common PD-linked mutations disrupt both functions. Furthermore, these studies show that even partial loss of Parkin function significantly alters excitatory neurotransmission and plasticity, suggesting that heterozygous *PARK2* mutations also impair these processes.

Although multiple studies show that Parkin regulates glutamatergic neurotransmission, it is currently unclear how its loss-of-function impacts excitatory drive in the brain. Several studies report that Parkin mutation/loss-of-function increases AMPAR and kainate receptor (KAR)-mediated currents in neurons, enhances glutamate excitotoxicity, and increases KAR and NMDAR levels in several brain regions (14, 17, 26, 54). These findings have led to the hypothesis that Parkin loss-of-function increases postsynaptic glutamate receptor levels and currents, promoting the death of dopaminergic neurons by excitotoxicity (14, 54). However, other groups report that Parkin mutation/loss-of-function decreases neuronal excitability and/or AMPAR EPSCs, as well as AMPAR levels in brain tissue from knockout mice and human patients (15, 16, 19, 26, 54, 55). We similarly find that Parkin loss-of-function decreases AMPAR-and NMDAR-mediated currents and cell-surface levels, suggesting that pathogenic Parkin mutations may reduce basal excitatory drive. On the other hand, our finding that Parkin deficiency/loss-of-function prevents the induction of LTD in hippocampal neurons, potentially leading to an inability to weaken or depotentiate glutamatergic synapses, lends support to the concept of increased excitation or altered excitatory/inhibitory balance in brains lacking Parkin.

Hippocampal NMDAR-dependent LTP is reportedly impaired in Parkin KO mice (20), and we find that cell-surface NMDAR levels are significantly reduced in neurons lacking enzymatically active Parkin due to its knockdown/KO or mutation, suggesting that synaptic plasticity mechanisms could be broadly disrupted. However, we find that LTP induction is intact in hippocampal neurons lacking functional Parkin, but that LTD is impaired. These findings suggest that the decrease in surface NMDARs does not disrupt all NMDAR-dependent plasticity mechanisms, although it may lead to more subtle deficits in LTP that we cannot detect in our analyses. These findings are also consistent with our previous work showing that Parkin is not required for AMPAR exocytosis, but is an important regulator of AMPAR endocytosis through its stabilization of Homer1-linked endocytic zones (19). Interestingly, none of the four Parkin mutants evaluated in this study were able to rescue either postsynaptic Homer1 levels or LTD induction, and all exhibited significantly reduced Homer1 binding capacity. These findings suggest that Parkin’s ability to mediate AMPAR internalization during LTD induction is intertwined with its ability to interact with and maintain Homer1 at the PSD.

*PARK2*-linked PD is often considered a distinct disease from idiopathic or other genetic forms of PD, with earlier onset, slower progression, and more purely motoric symptoms (7, 56). Consistent with these clinical phenotypes, *PARK2* patient brains lack widespread pathology, including the Lewy bodies found in other forms of PD, and cell loss is typically restricted to dopaminergic neurons of the substantia nigra (7). However, *PARK2* patients and heterozygous *PARK2* mutation carriers are reported to exhibit higher rates of neuropsychiatric symptoms (*e.g.* depression, psychosis, panic attacks) than non-carriers (7, 56, 57), and there is a significant association between *PARK2* heterozygosity and obsessive-compulsive disorder (58). While these disorders can be associated with altered dopaminergic signaling, they are also linked to changes in glutamatergic neurotransmission, including decreased AMPAR and NMDAR levels in specific brain regions (59-62). *PARK2* patients also exhibit unusual sensitivity to L-dopa treatment and rapid development of L-dopa-induced dyskinesia. Interestingly, dyskinesia in animal models and PD patients is linked to impaired glutamatergic synaptic plasticity, and in particular to the inability of synapses to become depotentiated/depressed following L-dopa induced potentiation (63-65). Our findings that Parkin-deficient excitatory synapses can become potentiated but not depressed, and that pathogenic Parkin mutations similarly do not support the induction of NMDAR-dependent LTD, may provide an explanation for the development of dyskinesia in *PARK2* patients. Moreover, increased potentiation of excitatory synapses would lead to higher levels of cell-surface AMPARs, potentially hastening the death of already-vulnerable dopaminergic neurons by glutamate excitotoxicity and thereby contributing to the early onset of motor symptoms in *PARK2* patients.

Together, our studies implicate glutamatergic synaptic dysfunction in the etiology of motor and non-motor symptoms experienced by *PARK2* patients. Intriguingly, recent genome-wide association studies report that *PARK2* deletions and copy number variations (CNVs) are also associated with autism, schizophrenia, and intellectual disability (35, 66-68), disorders linked to dysfunction of excitatory glutamatergic signaling. These new findings further underscore the importance of elucidating Parkin’s molecular mechanisms of action at glutamatergic synapses, both to facilitate the development of better treatments for motor and non-motor symptoms of *PARK2*-linked PD, and to shed light on its emerging role in the etiology of neuropsychiatric disease.

## Methods

### Antibodies and reagents

The following primary antibodies and dilutions were used for western blot and immunoprecipitation: mouse Parkin (Prk8, 1:1000; Santa Cruz Biotechnology), GluN1 (1:500; EMD Millipore), GluN2A (1:500; EMD Millipore), GluN2B (1:500; Neuromab), GluA1 phospho-Serine 845 rabbit antibody (1:1,000; Cell Signaling Technology), mouse tubulin (1:10,000; Sigma), rabbit tubulin (1:10,000; Abcam), rabbit GFP (1:1000; Invitrogen), mouse Myc (1:500; Santa Cruz Biotechnology), mouse HA (1:500; Santa Cruz Biotechnology), mouse Flag M2 (1:5,000; Sigma). DyLight fluorescent secondary antibodies were purchased from Thermo Fisher Scientific and diluted 1:15,000. The following primary antibodies and dilutions were used for immunostaining: purified human anti-GluN1 antiserum (a kind gift from C. Garner, German Center for Neurodegenerative Diseases (DZNE)) with 1 ug/ml working concentration, mouse GluA1 (1:100; EMD Millipore), rabbit Homer1 (1:500; Synaptic System), unconjugated anti-mouse/human secondary antibody (1:100; Invitrogen), mouse VAMP2 (1:500; Synaptic System), Alexa Fluor 568/647 anti-mouse/human secondary antibody (1:400; Invitrogen). Pharmacological agents used are as follows: cycloheximide (Calbiochem, 0.05 μg/μl, 24 h), NMDA, AP5, CNQX, and TTX (Tocris Bioscience), bicuculline, strychnine, leupeptin (50 μM), chloroquine (50 μM) and epoxomicin (0.1 μM)(Sigma), and PR-619 (LifeSensors, 50 μM). Unless otherwise indicated, all other chemicals are from Sigma-Aldrich.

### Primary hippocampal culture

Rat primary hippocampal cultures were prepared using a modified Banker culture protocol (Banker and Goslin, 1998; Waites et al., 2009). Briefly, neurons from embryonic (E18-E19) Sprague Dawley rat hippocampi, taken from animals of both sexes, were dissociated in TrypLE Express (ThermoFisher) for 20 min, washed with HBSS (Sigma), and plated in Neurobasal medium with B27 supplement and Glutamax (all from ThermoFisher) at a density of 250,000 neurons per well (12-well plates) or coverslip (22 × 22 mm square).

### Plasmids and transfection/transduction

The target sequence of shParkin and subcloning of shParkin +/-human Parkin constructs into pFUGW H1 have been described previously (19). The following plasmids were purchased from Addgene: pCI-SEP-GluA1/-GluA2/-GluN1/-GluN2B (all from Dr. Robert Malinow), pEGFP-Parkin C431S/W403A (from Dr. Edward Fon), pRK5-Myc-Parkin and pRK5-HA-ubiquitin (from Dr. Ted Dawson). Parkin mutants (T240M, R275W, R334C, G430D) were synthesized from Genewiz and subcloned into pEGFP-C2 and pFUGW H1-shParkin vector. pEGFP-Parkin C431S/W403A were also subcloned into FUGW H1-shParkin vector. Flag-GluN1 construct was a gift from O. Jeyifous (University of Chicago). The GFP-Homer1 construct was a gift from A. M. Grabrucker (University of Ulm, Ulm, Germany) and C. Garner. pFUGW constructs were used to generate lentivirus for transduction of primary neurons as described previously (33), except that Calfectin (SignaGen Laboratories) was used for transfection of HEK293T cells. HEK medium was replaced with Neurobasal medium 18–24 h after transfection, and this medium (viral supernatant) was harvested 24 h later. Neurons were transduced with 50–100 μl of lentiviral supernatant per well at 2–3 d *in vitro* (DIV) and used for experiments between 13-15 DIV. This time course was optimized based on efficacy of the shRNA knockdown. Transfections were performed on 5 DIV using Lipofectamine 2000 (Invitrogen) as described previously (19).

### Electrophysiology

Whole-cell patch-clamp recordings were performed in hippocampal neurons plated onto astrocyte microislands or 12mm coverslips and transduced with the indicated constructs, as previously described (19). Cultures were perfused at room temperature with an extracellular solution (145 mM NaCl, 5 mM KCl, 10 mM HEPES, 1.3 mM MgCl_2_, 2 mM CaCl_2_, 10 mM glucose) containing 1 μM of TTX (mEPSCs only) and 100 μM picrotoxin. Electrodes had resistances between 3-7MΩ containing intracellular solution (110 mm Cs-methanesulfonate, 10 mM Na-methanesulfonate, 10 mM EGTA, 1 mM CaCl_2,_ 10 mM HEPES, 10 mM TEA, 5 mM QX-314, 5 mM MgATP, 0.5 mM NaGTP). Neurons were visualized by fluorescence microscopy, and those expressing GFP were selected for recordings. For AMPA receptor current events, neurons were held at −65 mV and mEPSCs were recorded over 5 min using a MultiClamp 700B amplifier (Molecular Devices) controlled with a PC running MultiClamp Commander and pClamp (Molecular Devices) and pass filtered at 2 kHz. For miniature NMDA currents (NMDAR-EPSCs), neurons were voltage clamped at +40 mV and recorded over 5 min. For pharmacological induction of NMDARs, currents were measured following local application of NMDA (100 μM; Tocris Bioscience), using a custom-built application system ~200 μm from synapsing neurons on a specific microisland, in addition to bath application of the AMPAR antagonist CNQX (10 μM; Tocris Bioscience) while holding the neuron at +40 mV. Data acquisition and offline analysis of EPSCs were performed with pClamp (Clampex version 10.4). Approximately 15-20 neurons were analyzed for each experimental condition from three to four independent cultures.

### Immunoblotting

Cultured neurons were collected directly in 2x SDS sample buffer (Bio-Rad). Samples were subjected to SDS-PAGE, transferred to nitrocellulose membranes, probed with primary antibodies in 5% BSA/PBS plus 0.05% Tween 20 overnight at 4°C or 1 h at room temperature, followed by Dylight fluorescent secondary antibodies for 1 h. Membranes were imaged using an Odyssey Infrared Imager (model 9120; LI-COR Biosciences). Protein bands were quantified using the ImageJ (NIH) “Gels” function, and all bands were normalized to their loading controls.

### Biotinylation assay

Cultured cortical neurons transduced with GFP/shParkin/shParkin-WT (2-14 DIV) were washed with ice-cold PBS containing 1 mM MgCl_2_ and 0.1 mM CaCl_2_ (PBS+) and incubated with 1 mg/ml EZ-Link Sulfo-NHS-LC-biotin (ThermoFisher) in PBS+ for 20 min at 4°C with gentle agitation. Cells were washed with ice-cold quenching buffer (50 mM glycine in PBS+) for 10 min at 4°C, lysed in ice cold lysis buffer 50 mM Tris-base, 150 mM NaCl, 2% Triton X-100, 0.5% deoxycholic acid, and supernatant was incubated with streptavidin-Sepharose beads (ThermoFisher) for 2 h at 4°C. Bound proteins were immunoblotted with GluN1, GluN2A, GluN2B, Parkin and tubulin. The data were quantified by measuring surface receptor to total input receptor band intensity ratios using ImageJ software and normalizing to GFP control cultures.

### Co-immunoprecipitation assay

For co-immunoprecipitation studies, HEK293T cells were transfected with plasmids (pCI-SEP-GluA1/-GluA2/-GluN1/-GluN2B and Myc-Parkin for Myc-IP assay; GFP-Homer1 and GFP control/GFP-Parkin WT/T240M/R275W/R334C/G430D for Homer1 IP assay) using Calfectin according to the manufacturer’s (SignaGen Laboratories) protocol. Cell lysates were collected 36 h after transfection in lysis buffer (50 mM Tris-base, 150 mM NaCl, 1% Triton X-100, 0.5% deoxycholic acid) with protease inhibitor mixture (Roche) and clarified by centrifugation at high speed (20,000 rcf). The resulting supernatant was incubated with Dynabeads (ThermoFisher) coupled with anti-Myc or anti-Homer1 antibodies at 4°C under constant rotation for 2-4 h. Beads were washed two to three times with lysis buffer and once with PBS. Bound proteins were eluted using sample buffer (Bio-Rad) and subject to SDS-PAGE immunoblotting.

### Ubiquitination assay

HEK293T cells were transfected with pCI-SEP-GluA1/-GluA2/-GluN1/-GluN2B, Myc-Parkin and HA-ubiquitin, or with GFP/GFP-Parkin WT/-Parkin C431S/-Parkin W403A/-Parkin T240M/-Parkin R275W/-Parkin R334C/-Parkin G430D, Flag-GluN1 and HA-Ub using Calfectin according to the manufacturer’s protocol. 36 h after transfection, proteasome and lysosome inhibitors including leupeptin, chloroquine, epoxomicin, and PR619 were added to the cultures. After 2-4 h treatment, cell lysates were collected in lysis buffer with protease inhibitor mixture and clarified by centrifugation at high speed (20,000 rcf). The resulting supernatant was incubated with Dynabeads (ThermoFisher) coupled with anti-GFP antibodies or anti-Flag M2 magnetic beads (Sigma) at 4°C under constant rotation for overnight. Beads were washed two to three times with lysis buffer and once with PBS. Bound proteins were eluted using sample buffer (Bio-Rad) and subject to SDS-PAGE immunoblotting.

### Immunofluorescence microscopy

Primary antibodies and concentrations are listed above. Alexa Fluor 488-, Alexa Fluor 568-, or Alexa Fluor 647-conjugated secondary antibodies (ThermoFisher) were used at 1:400. Neurons were immunostained as described previously (33). Briefly, coverslips were fixed with Lorene’s fixative (60 mM PIPES, 25 mM HEPES, 10 mM EGTA, 2 mM MgCl2, 0.12 M sucrose, 4% formaldehyde) for 15 min, primary and secondary antibody incubations were performed in blocking buffer (2% glycine, 2% BSA, 0.2% gelatin, and 50 mM NH4Cl in 1× PBS) for 1–2 h at room temperature or overnight at 4°C, and all washes were done with PBS. For surface labeling, GluN1 (1 μg/ml) or GluA1 antibody (1:100) was added to live neurons, incubated for 30 min at room temperature, washed three times in PBS, and fixed and stained with secondary antibodies as described. Coverslips were mounted with VectaShield (Vector Laboratories) and sealed with clear nail polish. Images were acquired with a 63× objective (Neofluar, NA 1.4) on a Zeiss LSM 800 confocal microscope running Zen2 software.

### GluN1 internalization assay

GluN1 internalization was measured using a protocol as described previously (19). Neurons expressing GFP/shParkin/shParkin-WT were incubated with human GluN1 antibody (1 μg/ml in PBS) for 30 min at room temperature to label surface NMDARs, washed four times with PBS, and either fixed immediately with Lorene’s fixative or incubated at 37°C for 15 min before fixation to allow for receptor internalization. Cells were then incubated with anti-human Alexa Fluor 647 secondary antibody diluted in blocking buffer (used for all remaining steps) for 40 min at room temperature, washed three times, and incubated for 30 min with unconjugated anti-mouse secondary (1:100) antibody to block any remaining unlabeled cell-surface GluN1 antibody. Finally, cells were washed four times, permeabilized with 0.25% Triton X-100, and incubated for 1 h at room temperature with anti-human Alexa Fluor 568 to label internalized GluN1.

### GluN1 recycling assay

Neurons were incubated with anti-GluN1 antibody to label surface receptors as described above and incubated at 37°C for 30 min to allow for receptor internalization. All non-internalized surface antibody was subsequently blocked with an unconjugated anti-human secondary antibody at room temperature for 30 min and washed four times in PBS. Next, neurons were fixed or incubated at 37°C for 1 h to allow for receptor recycling back to the plasma membrane. They were then fixed, incubated for 40 min with Alexa Fluor 568-conjugated anti-mouse secondary antibody diluted in blocking buffer (used for all remaining steps) to label the recycled surface population of receptors, and washed three times. The remaining surface primary antibodies were blocked with an excess of an unconjugated anti-human secondary antibody at room temperature for 30 min, and neurons again were washed four times. After permeabilization, neurons were incubated with Alexa Fluor 647-conjugated anti-human secondary antibody for 40 min to label the internalized population of receptors, washed three times, and mounted and imaged as described previously.

### Chemical LTP/LTD assay

Chemical LTP/LTD assay was performed as described previously (34). Neurons were washed twice in Tyrodes solution (120 mM NaCl, 2 mM CaCl_2_, 2.5 mM KCl, 25 mM HEPES, 30 mM glucose, pH 7.4), and then incubated in cLTP buffer (200 μM glycine, 1 μM strychnine, 20 μM bicuculline in Tyrodes solution, pH 7.4) for 5 min. Following a quick wash with wash buffer (2 mM MgCl_2_, 50 μM AP5, 10 μM CNQX, 0.5 μM TTX, 1 μM strychnine in Tyrodes solution, pH 7.4), neurons were incubated for another 5 min in this buffer prior to fixation. For chemical LTD, neurons were washed twice in Tyrodes solution, then incubated with cLTD buffer (25 μM NMDA, 1 μM strychnine, 0.5 μM TTX, 10 μM glycine in Tyrodes solution, pH 7.4) for 5 min followed by a quick wash with wash buffer. After cLTP or cLTD treatment, neurons were subjected to surface GluA1 staining for 10 min, then fixed and immunostained with secondary antibody.

### Image analyses

Images of surface or total GluN1/GluA1 were manually thresholded to select only puncta that were greater than 2-fold above image background. The average intensity values of puncta were measured using the “Analyze Particles” function in ImageJ/Fiji, with puncta size set between 2 and 100 pixels. Intensity of Homer1 puncta at sites of VAMP2 colocalization was calculated by creating a selection in ImageJ/Fiji using the VAMP2 channel, applying it to the relevant Homer1 channel, and measuring average intensity values of Homer1 puncta using the “Analyze Particles” function, with puncta size set between 2 and 100 pixels. For internalization and recycling assays, ratios of internalized GluN1 intensity to surface GluN1 intensity, or recycled to internalized GluN1 intensity, were calculated per punctum (after measuring raw intensity values in the red and far-red channels) and averaged for each field of view. These averaged ratios were then normalized to the control condition (average value set to 1) and expressed as a fraction of control.

### Statistical analysis

Statistical analyses were performed in GraphPad Prism5 software using either one-way ANOVA-Tukey, ANOVA-Dunnett’s multiple comparison test, or unpaired, two-tailed Student’s *t* tests, with p<0.05 considered significant. Data are presented as mean ±SEM.

## List of abbreviations

PD: Parkinson’s disease
AMPA: α-amino-3-hydroxy-5-methyl-4-isoxazolepropionic acid
NMDA: N-methyl-D-aspartic acid
GluA: AMPA receptor
GluN: NMDA receptor

## Authors’ contributions

M.Z., G.P.C. and C.L.W. designed the research; G.P.C. did the electrophysiology experiments and analyses; M.Z. performed all other experiments and data analyses; M.Z. and C.L.W. wrote the manuscript.

## Acknowledgements

We thank Dr. Craig Garner (German Center for Neurodegenerative Diseases (DZNE), Universitätsmedizin Berlin) for the generous gift of human GluN1 antibody, Drs. Okinola Jeyifous and William Green (University of Chicago) for pFlag-GluN1 construct, Dr. Robert Malinow for pCI-SEP-GluA1/-GluA2/-GluN1/-GluN2B constructs (Addgene plasmids #23997-24001), Dr. Edward Fon for pEGFP-Parkin WT/C431S/W403A constructs (Addgene plasmids #45875-77), Dr. Ted Dawson for pRK5-Myc-Parkin and pRK5-HA-Ubiquitin-WT constructs (Addgene plasmids #17612 and #17608, respectively), and Dr. A.M. Grabrucker (University of Ulm, Germany) for GFP-Homer1 construct. We would also like to thank Dr. Damian Williams (CUMC electrophysiology core) for technical help with electrophysiology experiments, and Cyndel Vollmer for help with glial microisland preparation.

